# MEGA12: Molecular Evolutionary Genetic Analysis version 12 for adaptive and green computing

**DOI:** 10.1101/2024.12.10.627672

**Authors:** Sudhir Kumar, Glen Stecher, Michael Suleski, Maxwell Sanderford, Sudip Sharma, Koichiro Tamura

## Abstract

We introduce the 12th version of the Molecular Evolutionary Genetics Analysis (*MEGA*) software. This latest version brings many significant improvements by reducing the computational time needed for selecting optimal substitution models and conducting bootstrap tests on phylogenies using maximum likelihood (ML) methods. These improvements are achieved by implementing heuristics that minimize likely unnecessary computations. Analyses of empirical and simulated datasets show substantial time savings by using these heuristics without compromising the accuracy of results. *MEGA12* also implements an evolutionary sparse learning approach to identify fragile clades and associated sequences in evolutionary trees inferred through phylogenomic analyses. In addition, this version includes fine-grained parallelization for ML analyses, support for high-resolution monitors, and an enhanced *Tree Explorer*. The *MEGA12* beta version can be downloaded from https://www.megasoftware.net/beta_download.

## INTRODUCTION

The Molecular Evolutionary Genetics Analysis (*MEGA*) software is extensively used for molecular evolution and phylogenetics (Kumar 2022). It offers many computational tools, including Maximum Likelihood (ML), Maximum Parsimony (MP), Ordinary Least Squares (OLS), Bayesian, and distance-based methods (**Fig. 1**). Some popular functionalities in *MEGA* are the selection of optimal nucleotide and amino acid substitution models, inference of evolutionary relationships, tests of phylogenies using the bootstrap method, estimation of sequence divergences and times, and the reconstruction of ancestral sequences (***Supplementary Fig. S1***).

**Figure 1.**
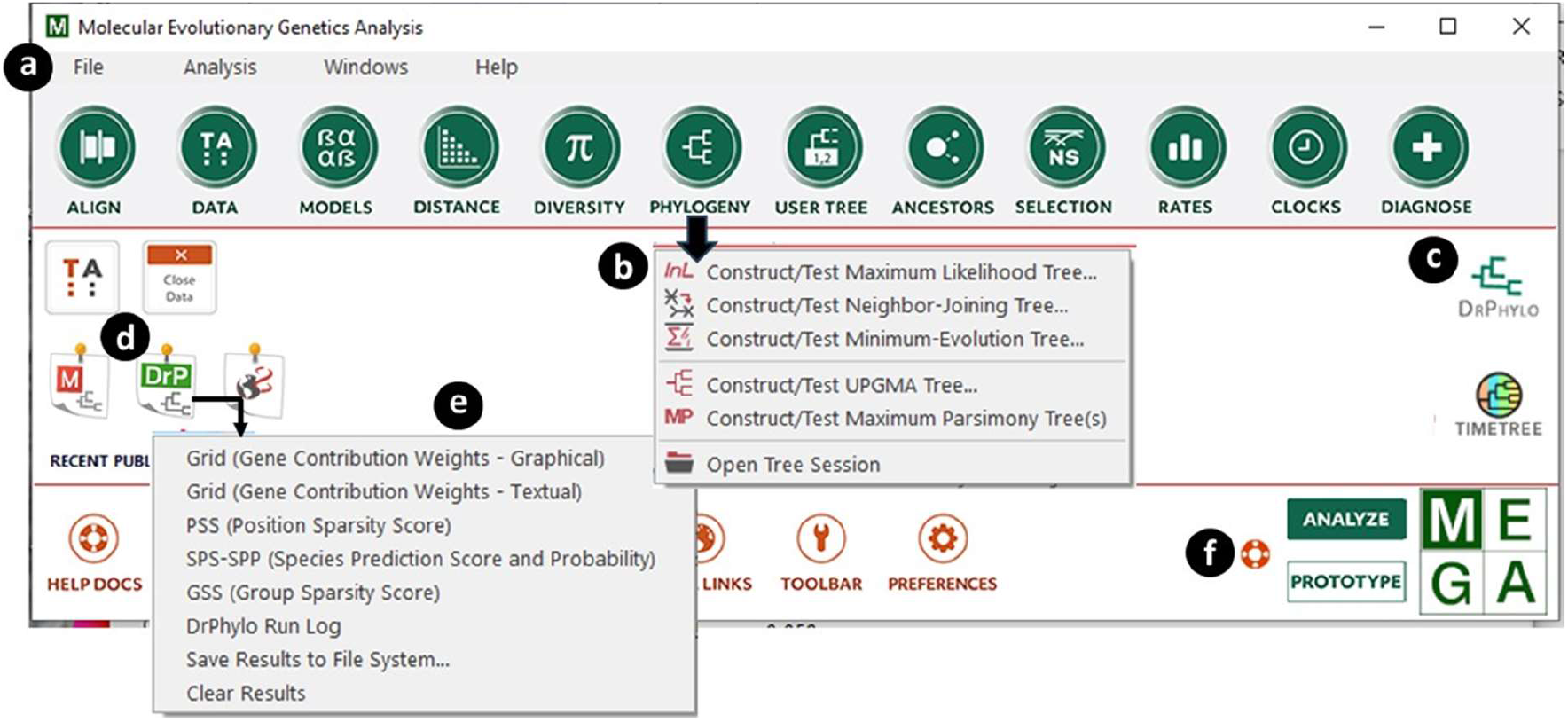
Main Graphical User Interface (GUI) of *MEGA12*. (***a***) The main toolbar provides access to various analytical capabilities organized in drop-down menus. (***b***) One of the drop-down menus is shown on the main window. (***c***) The *AppTile* provides access to the linked *DrPhylo* application, which is also accessible from the *Tree Explorer* window (see **Fig. 4a**). (***d***) The *OutputTile* provides access to results from *DrPhylo* analysis via a drop-down menu (***e***). (***f***) Clicking the *Prototype* button allows for the building of a ‘mao’ analysis configuration file for the command line analysis using MEGA-CC. It is necessary to click *Analyze* to return to the standard mode to conduct analysis using the GUI.

Notably, phylogenetic ML analyses are time-consuming and contribute to a substantial carbon footprint (Kumar 2022). To tackle these challenges, the latest update of *MEGA* has aimed to improve the computational efficiency of ML analyses. This has been achieved by developing and implementing heuristic approaches that avoid unnecessary calculations while preserving accuracy. Further efficiencies are gained by optimizing parallel computations to make the best use of available computing resources. In the following sections, we will discuss these updates and enhancements to the Graphic User Interface (GUI), including seamless access to an external application (*DrPhylo*) for detecting fragile clades and associated sequences in the inferred phylogenies.

## RESULTS

### Adaptive computing in selecting the optimal substitution model

Selecting the most suitable substitution model is often the initial step in molecular phylogenetics. The ML method for model selection was initially introduced in *MEGA5* (Tamura et al. 2011) and has been frequently used (***Supplementary Figure S1***). *MEGA* assesses six primary nucleotide substitution models to determine the optimal model: General Time Reversible (GTR), Hasegawa-Kishino-Yano (HKY), Tamura-Nei (TN93), Tamura 3-parameter (T92), Kimura 2-parameter (K2P), and Jukes-Cantor (JC); see (Nei and Kumar 2000) for a review. These primary substitution models describe the instantaneous probabilities of nucleotide substitutions at individual sites. They can be combined with a (discretized) Gamma distribution of rate variation among sites (indicated by +G) and the presence/absence of invariant sites (indicated by +I), which are reviewed in Nei and Kumar (2000).

The computational time required for the ML analysis of 24 different model combinations increases with data size (**Fig. 2a**). Here, data size is quantified by multiplying the number of sequences (S) by the distinct number of site configurations (C) in the multiple sequence alignment (MSA). Distinct site configurations are used because all sites with the same configuration, patterns of bases present across sequences, are compressed into a single column with a corresponding frequency in the ML analysis (Sharma and Kumar 2021). Due to the extensive use of model selection by a large user base of *MEGA* (***Supplementary Figure S2***), these analyses’ combined computational cost – and consequently, energy consumption-is enormous.

**Figure 2.**
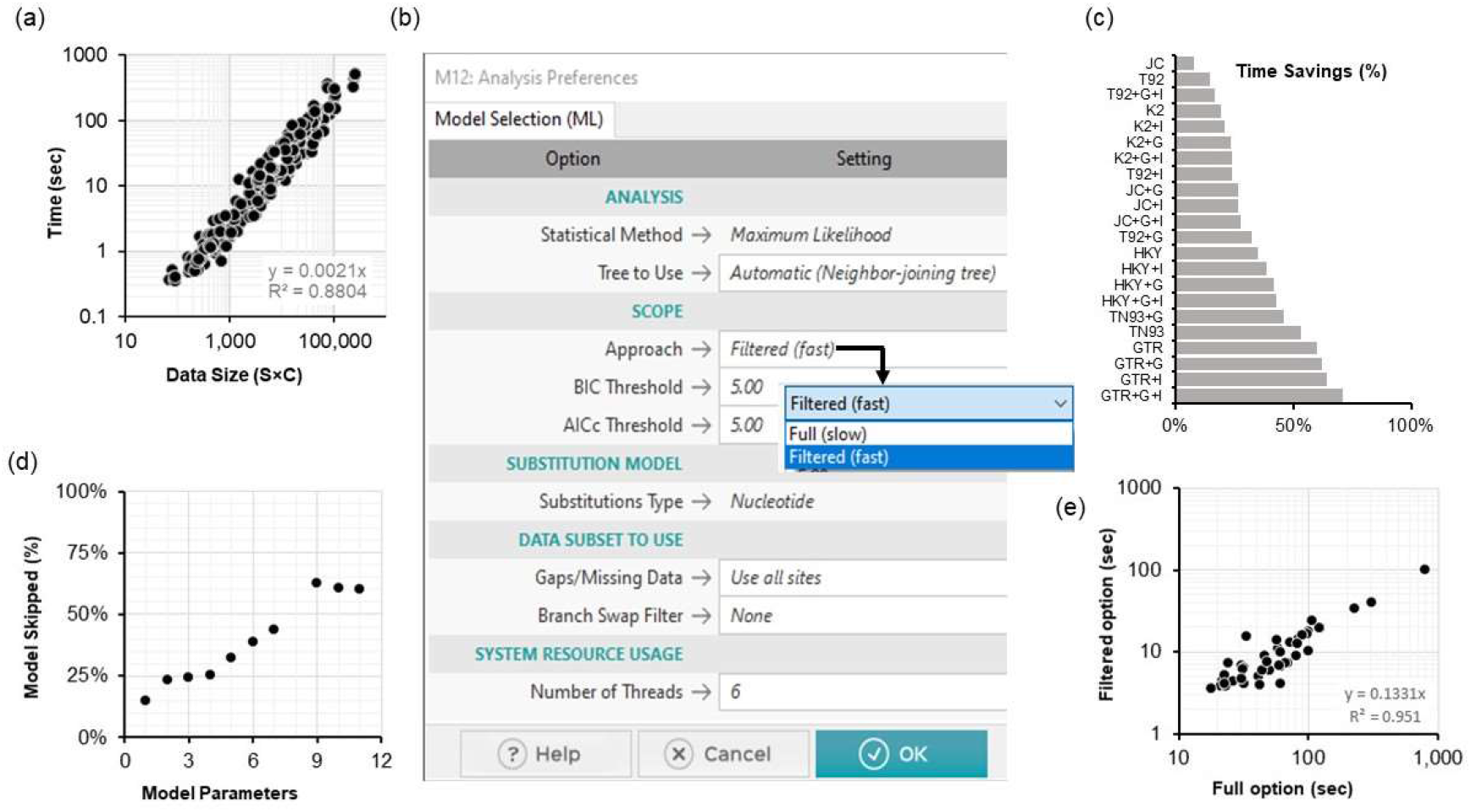
Substitution model selection using *MEGA12*. (***a***) The relationships between the time required for the standard model selection and the data size: the product of the number of sequences (S) and the number of distinct site configurations (C) in the sequence alignment. The time required for model selection analysis increases linearly with the data size. (***b***) *MEGA12*’s *Analysis Preferences* dialog box allows users to set options for model selection analysis. The newly added *Filtered* option is shown, which offers a setting of BIC and AICc thresholds. As the main text explains, a smaller number will result in testing fewer models. (***c***) Time savings are achieved using the *Filtered* option with default parameters, which is the greatest for datasets for which the full analysis selects a complex best-fit substitution model. (***d***) The relationship between the number of model parameters and the average percentage of model combinations whose ML evaluation was skipped. (***e***) The relationship of time taken with the *Filtered* and *Full* options for model selection for chloroplast amino acid MSAs. The slope of the regression line is 0.18, indicating that the *Filtered* option greatly speeds model selection.

To accelerate model selection, we have developed a heuristic approach that reduces the number of substitution model combinations tested, eliminating potentially suboptimal models early in the process. This new heuristic for model selection analysis of nucleotide sequence alignments begins with evaluating the ML model fit for six base models: GTR, HKY, TN93, T92, K2P, and JC. MEGA uses the Bayesian Information Criterion (BIC) and the corrected Akaike Information Criterion (AICc) to evaluate model fit where these criteria are calculated based on the log-likelihood fit of each model to the given multiple sequence alignment (MSA) and its associated parameters (Tamura et al., 2011). Therefore, MEGA first determines the BIC and AICc for these six base models. The base model with the lowest BIC value (BIC_min_) is selected first, and AICc_min_ is taken as the AICc value of this same model. Base models with BIC or AICc values not exceeding 5 points of BIC_min_ or AICc_min_, respectively, are considered potentially optimal models for further consideration. Models with AICc or BIC exceeding 5 points of AICc_min_ or BIC_min_ are considered sub-optimal, and other model combinations derived from these suboptimal base models are not tested further.

The ML analysis of the remaining base models in combination with +I, +G (with four categories), and both +I+G is then carried out to calculate different information criteria for the final selection. The model with the lowest BIC is determined as the best-fit model of substitutions. MEGA outputs different information criteria scores, log likelihood (*lnL*), and other model parameters for models tested. In *MEGA12*, this heuristic can be used by selecting the newly added *Filtered* option (**Fig. 2b**). Users can also choose to set a desired threshold for BIC and AICc, with smaller threshold values resulting in the elimination of more models earlier.

In an analysis of 240 simulated datasets generated with substitution models of various complexities in an independent study (Abadi et al. 2019) (see ***Materials and Methods***), the *Filtered* option identified the same optimal substitution model as the *Full* analysis for 240 datasets (100% concordance). It achieved as much as 70% reduction in computational time when a complex substitution model fits the best (see **Fig. 2c**). The savings were lower for simpler models because simpler models are nested within more complex models, so the latter may not be eliminated early on. For this reason, the number of model combinations analyzed is directly related to the number of parameters in the best-fit model (see **Fig. 2d**). We anticipate that most users will experience substantial speed-ups by utilizing the *Filtered* option because complex models are often the optimal fit for bigger datasets.

The *Filtered* option is also implemented for amino acid MSAs. In this case, *MEGA12* first determines BIC and AICc in all eight primary substitution models and eliminates all models with BIC and AICc values five more than BIC_min_ or AICc_min_, respectively, as outlined above. In the next step, BIC and AICc are computed for each of the remaining models combined with the +F option, using the empirical frequencies from the MSA (Tamura et al. 2011). New model combinations are eliminated if their BIC and AICc exceed BIC_min_ or AICc_min_, respectively, by five or more. In the final step, the ML analysis of the remaining models in combination with +I, +G, and +G+I is then carried out to generate the final result.

The use of the *Filtered* option for 45 chloroplast proteins (31-1388 amino acids) from 10 species resulted in extensive (∼87%) savings compared to the *Full* analysis (see **Fig. 2e**). The same best-fit substitution models were found as the *Full* option for 43 proteins (95% concordance). A statistically indistinguishable model (ΔBIC < 10) was chosen for one of the other two datasets, while the second-best model was selected for the other. Model selection with the *Filtered* option for concatenating 45 chloroplast protein MSAs (11,039 amino acids) took only 3.67 minutes instead of 27.12 minutes for the *Full* analysis while producing the same substitution model.

These results suggest that the speed-up for model selection analysis will be realized for every dataset analyzed with the filtered option in *MEGA12*. However, the degree of computational efficiency gained depends on the complexity of the substitution model that best fits the data.

### Adaptive Bootstrapping

After selecting the optimal substitution model, the next step in phylogenetic analysis is to infer evolutionary relationships and assess the confidence in the monophyly of inferred clades. The bootstrap approach has been available in *MEGA* since version 1 to estimate confidence in the inferred relationships (Felsenstein 1985; Kumar et al. 1994). In the bootstrap procedure, many resampled MSA are generated by sampling sites with replacement until the number of sites in the resampled MSA is the same as in the original MSA. Phylogenies are inferred from these resampled MSAs using a phylogenetic tree estimation approach (e.g., ML approach). The proportion of times a cluster of sequences appears in the phylogenies obtained from the resampled MSAs is its bootstrap support (BS). A high BS value indicates that the inferred clade is statistically supported (Felsenstein 1985).

Users frequently choose to generate a large number of resampled MSAs (500 - 2000 replicates) because every BS value is an estimate whose accuracy is determined by the number of resampled MSAs analyzed (Hedges 1992; Pattengale et al. 2010). The number of bootstrap replicates increases the time required proportionally, which becomes particularly onerous for computationally intensive ML phylogenetics. To reduce this burden, *MEGA12* introduces an *Adaptive* option for bootstrap analysis (**Fig. 3a**), automatically determining the optimal number of replicates for the bootstrap analysis. It is based on the fact that high BS values, which are of primary interest to researchers, can be estimated with high precision (i.e., low standard error [SE]) from a small number of replicates. For example, the estimation of BS = 95% with an SE = 2.5% requires only 75 bootstrap replicates. Interestingly, BS values close to 50%, often of limited biological interest, require hundreds of replicates (390) to reach an SE of 2.5%. Therefore, using a large number of replicates primarily increases the precision of BS values close to 50%. To emphasize that the BS values are estimates with standard errors, we have updated *Tree Explorer* in *MEGA12* to display the range of BS values (± 1 SE, **Fig. 3b**). Users have the option to display the BS value or the range calculated using the above formula. This display option is also available for phylogenies inferred using distance-based (e.g., Neighbor-Joining) and MP methods.

**Figure 3.**
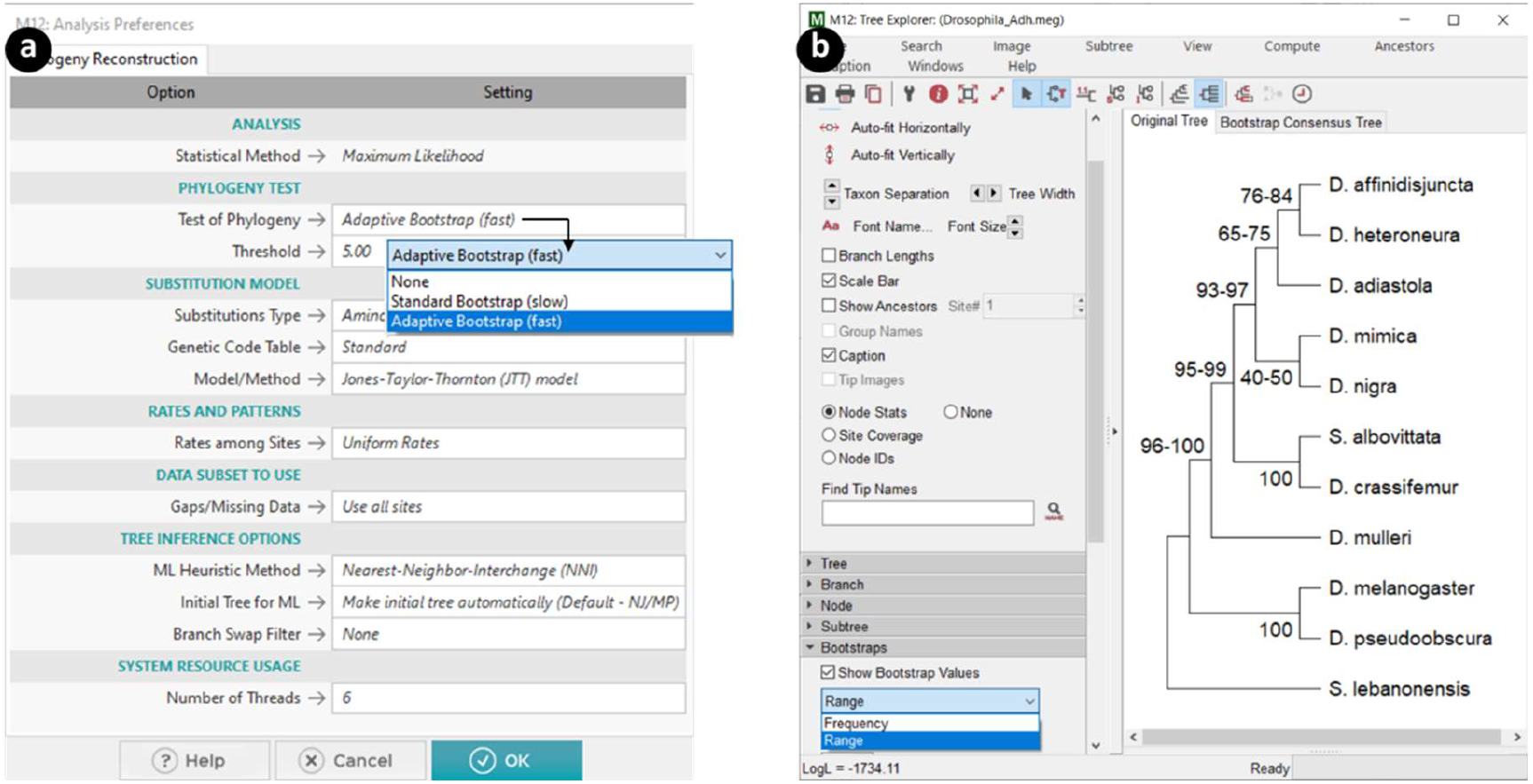
Adaptive bootstrap analysis of the Drosophila *Adh* dataset. (***a***) *MEGA12*’s *Analysis Preferences* dialog box with the new *Adaptive* option for automatically determining the number of bootstrap replicates. The use of the *Adaptive* option stops generating bootstrap replicates when the standard error (SE) of every bootstrap support (BS) value in the phylogeny becomes less than 5% (default option). The *Threshold* option allows setting an alternative SE value for more or less precise BS values. (***b***) *MEGA*’s updated *Tree Explorer* with an option to display the range of BS values (± 1 SE) for every inferred clade. The results are based on the analysis of the data in the *Drosophia_Adh*.*meg* file distributed in the *Examples* folder with *MEGA12*.

*MEGA12*’s *Adaptive* option first generates 25 resampled MSAs. If all the BS values in the phylogeny have an SE < 5%, then additional resampled MSAs are generated until all the BS values in the phylogeny have achieved an SE < 5%, a threshold that can be set by the user (**Fig. 3a**). Our rationale for picking a 5% default was that some of the clades in the inferred phylogeny would have BS close to 50%, so achieving an SE < 5% for these clades will require many replicates (often ∼100). With 100 replicates, the clades with high BS values will have SE closer to 2.5%. This is evident from the analysis of *Drosophia_Adh*.*meg* dataset distributed in the *Examples* folder with *MEGA12* (**Fig. 3b**). *MEGA12* stopped the bootstrap procedure after 87 replicates, which resulted in much narrower ranges (BS-SE to BS+SE) for high BS values than low BS values (**Fig. 3b**).

We also analyzed all 240 simulated datasets with standard and adaptive bootstrap approaches. **Figure 4a** shows the relationship of the BS values obtained using 500 replicates (x-axis) and determined adaptively (y-axis) for 240 datasets. The relationship is strong (R^2^ = 0.99) with a slope close to 1 (**Fig. 4a**), i.e., the two approaches produced very similar results overall. However, there is some dispersion because we used a threshold of SE = 5% for adaptive bootstrapping. The number of replicates needed for adaptive bootstrapping varied between 25 and 124 for the 240 datasets, with 25 because of a hard lower limit placed by *MEGA12*. Because the default SE threshold of 5% is applied to every clade in the inferred phylogeny, the number of bootstrap replicates needed will be determined by the node whose estimated BS support is closest to 50% because the variance of a BS value (say *b*) is given by *b*(1- *b*)/*r*, where *r* is the number of replicates (Hedges 1992). Indeed, there is a direct relationship between the number of replicates needed in adaptive bootstrapping and the minimum |BS - 50%| value in the inferred phylogeny in the analysis of 240 datasets (**Fig. 4b**). For phylogenies with uniformly high BS values (bottom right in **Fig. 4b**), the number of replicates needed is much smaller than when the phylogeny contains even one BS value close to 50%. In any case, the *Adaptive* option resulted in speed-ups on an average of 81% times (**Fig. 4c**), ranging from 61% to 95% for these datasets. In general, we expect such speed-ups to be realized for all datasets analyzed in MEGA, with actual computational efficiency depending on the properties of the data and the bootstrap stability of the phylogenetic inference.

**Figure 4.**
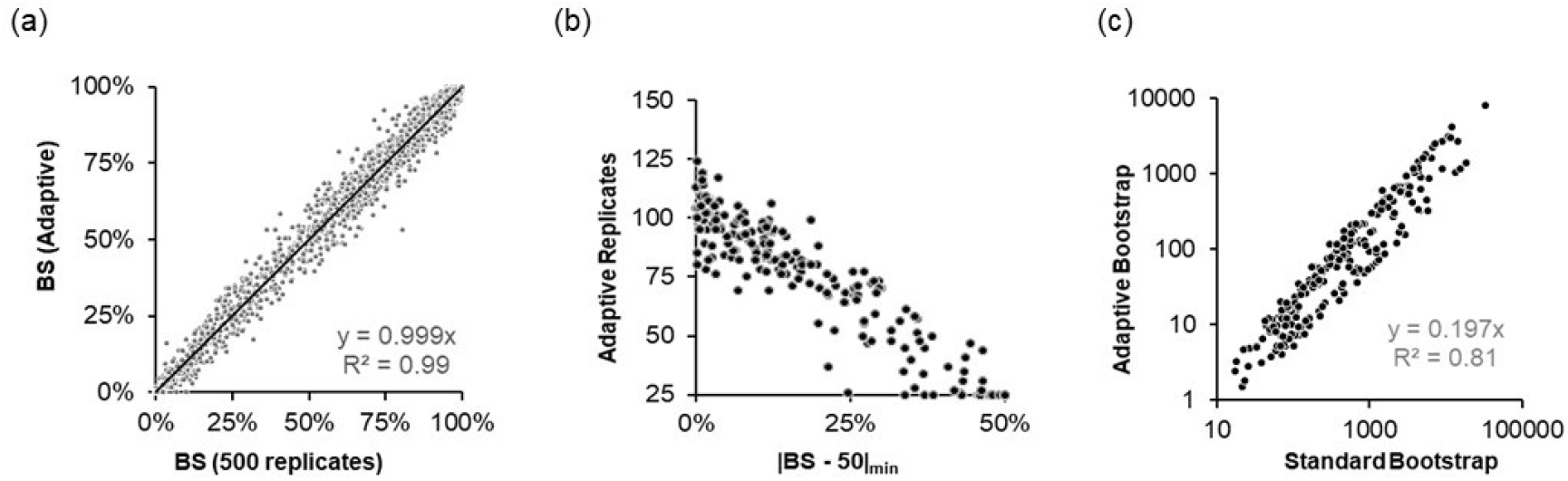
Adaptive bootstrap analysis in *MEGA12*. (***a***) Comparison of BS values obtained using the *Adaptive* determination of the number of bootstrap replicates (y-axis) and those obtained using 500 bootstrap replicates (x-axis). Results from all 240 data sets were pooled together. The slope of the linear regression through the origin is 0.99 (*R*^2^ = 0.99). (***b***) The relationship between the minimum |BS - 50%| in phylogeny and the number of replicates needed by the *Adaptive* analysis. The negative trend (correlation = -0.96) confirms the inverse relationship expected theoretically. (***c***) The relationship of time taken between the *Adaptive* and *Standard Bootstrap* approach for estimating statistical support for clade relationships inferred for simulated DNA sequence alignments. The slope of the regression line is ∼0.20, indicating that the *Adaptive* approach speeds up the bootstrap support estimation significantly.

### Integration of the *DrPhylo* application to assess the fragility of inferred clades

MEGA currently uses the concatenation supermatrix approach when the input contains multiple genes, domains, or genomic segments. This approach is effective in producing organismal relationships with high confidence (Gadagkar et al. 2005; Kumar, Filipski, et al. 2012; Song et al. 2012; Kapli et al. 2020; Williams et al. 2020; Sharma and Kumar 2021). However, the concatenation supermatrix approach may occasionally lead to incorrect or fragile inferences with very high bootstrap support due to systematic, modeling, and data-specific biases (Gatesy and Springer 2014; Warnow 2015; Sharma and Kumar 2024). *MEGA12* now makes available the *DrPhylo* approach, a sparse learning approach (Kumar and Sharma 2021; Sharma and Kumar 2024) to identify inferred clades that may be formed due to data-specific biases (Sharma and Kumar 2024). Users can launch *DrPhylo* analysis for any clade in the phylogeny displayed in the *Tree Explorer* (**Fig. 5a**), as well as directly from the main *MEGA* window (**Fig. 1b**). One can choose to use partitions (genes or segments) in the currently active dataset or divide the data into segments of equal length (**Fig. 5b**). The user may also provide a list of files, each containing a sequence alignment for a data segment, for *DrPhylo* analysis (**Fig. 5c**).

**Figure 5.**
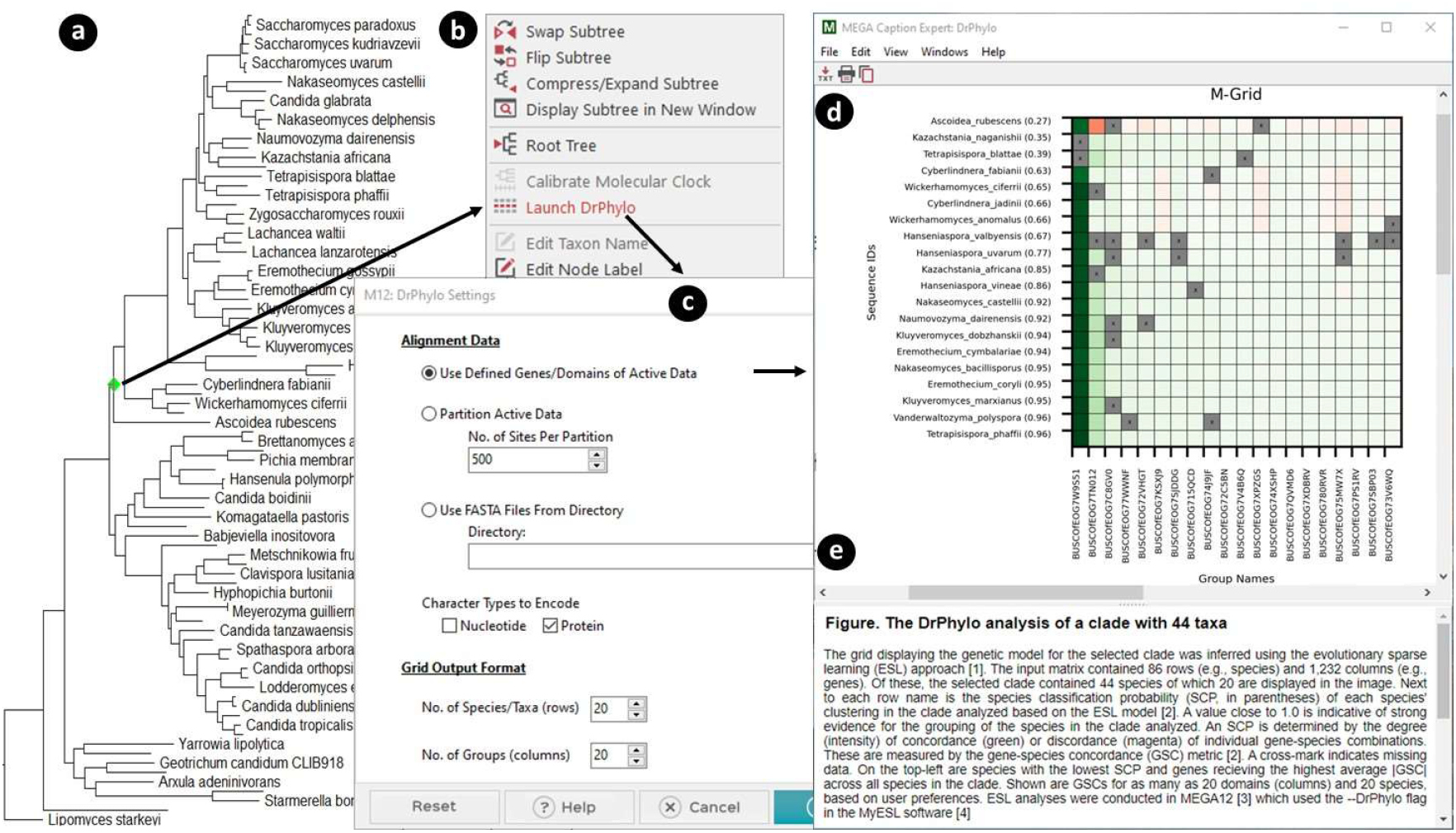
Conducting *DrPhylo* analysis via the *Tree Explorer* in *MEGA12*. (***a***) Users select the clade of interest by clicking on its ancestral branch or node (highlighted in green) in the *Tree Explorer* window. (***b***) The context-sensitive menu, which includes the *Launch DrPhylo* option, is displayed. (***c***) The dialog box to make selections for *DrPhylo* analysis. (***d***) A graphical representation of the genetic model of the selected clade in a grid format (M-Grid) is shown along with a descriptive caption. This model and other output files are accessible from the *DrP OutputTile* (see **Fig. 1c**). (***e***) Caption showing the details of the *DrPhylo* analyses and a description of the results.

*DrPhylo* is implemented in the *MyESL* software (Sanderford et al. 2024) and linked with *MEGA12* by upgrading the source code that has been previously used to link *Muscle* (Edgar 2004) with *MEGA3* (Kumar et al. 2004). *MEGA12* extracts the application binaries and resources to set up the *DrPhylo* execution environment with the proper files, structure, and permissions when *DrPhylo* is run in *MEGA12* for the first time after installation.

We demonstrate the steps of *DrPhylo* analysis and its outputs by analyzing a phylogenetic tree of 86 fungal species inferred from the ML analysis of a concatenated sequence alignment of 1,232 genes (Shen et al. 2017). In this phylogeny, the placement of *Ascoidea rubescens* in the ML phylogeny was reported to be contentious, even though it received 100% standard bootstrap support (Sharma and Kumar 2024; Shen et al. 2017). The clade of interest contains *A. rubescens* and 43 other species, which is easily disrupted by excluding just one gene (Shen et al. 2017). In *MEGA12*, the user selects the clade by clicking on the stem branch (**Fig. 5a**). Then, *DrPhylo* analysis is launched by clicking on the menu that appears (**Fig. 5b**) and choosing options to specify the sequence data, data type, and model grid size (**Fig. 5c**). We selected the dataset that was already active in MEGA and chose to display 20 top genes along with 20 taxa from the clade-of-interest (**Fig. 5c**).

*MEGA12* automatically prepares the input sources (alignments and the tree file) for a given analysis and copies them to *DrPhylo*’s execution environment. *DrPhylo* is launched in a separate child process along with command line arguments. As *DrPhylo* executes, *MEGA* captures all analysis-progress information and displays it in a new progress window. When the *DrPhylo* process finishes, *MEGA* loads all of the results files into memory and adds a *DrP OutputTile* (**Fig. 1d**) to the main form to provide access to output files produced by *DrPhylo* (**Fig. 1e**). Output files include model grid (M-grid) in both graphical and textual format, position and gene sparsity score (PSS and GSS), DrPhylo run log, and options for saving or erasing results; see (Sharma and Kumar 2024; Kumar and Sharma 2021) for more details. Clicking on a menu item in the *DrP OutputTile* smartly displays the result. For example, clicking the first menu item displays the graphics file in the integrated web browser with the figure legend (**Fig. 5d**).

In the grid displayed (called the model grid; M-grid), the horizontal axis represents the genes included in the model, while the vertical axis represents the species along with the classification probability for their membership within the clade of interest. The clade probability (CP) is the smallest probability of these probabilities because the most unstable member dictates the fragility of the clade. Each cell (say *i*,*j*) in the M-grid reflects the gene-species concordance (GSC) score, which quantifies the support a gene (*j*) provides for a species (*i*) within the clade of interest. Green cells indicate the gene’s positive support for species membership within the clade, whereas red cells display gene-species combinations that harbor a discordant signal. Based on the GSC score, the color intensity corresponds to the strength of the concordance or discordance. Sharma and Kumar (2024) provided a detailed explanation of the M-grid for this dataset.

The M-grid in **Fig. 5d** shows that the CP for the clade analyzed is 0.27 due to *A. rubescens* receiving a low membership probability. One gene, BUSCOfEOG7W9S51, provides the strongest support for the placement of *A. rubescens* (green cells) in this clade, but many other genes did not support (red) this placement. Interestingly, BUSCOfEOG7W9S51 also strongly supports other species’ inclusion inside the clade. Therefore, BUSCOfEOG7W9S51 is highly influential for this clade and places *A. rubescens* inside it. This gene was also found using the ML approach by comparing two alternative hypotheses for the placement of *A. rubescens* (Shen et al. 2017). Removing this gene from the analysis changed the placement of *A. rubescens* with moderate bootstrap support (Shen et al. 2017; Sharma and Kumar 2024). This is because of BUSCOfEOG7TN012, which shows a very high disagreement with the placement of *A. rubescens* in this clade (red cell, **Fig. 5d**) but not others. Such disagreement discordance was confirmed in the gene tree, which positioned *A. rubescens* far outside the clade of interest (Sharma and Kumar 2024). In summary, the availability of *DrPhylo* in *MEGA12* will allow users to investigate fragile species relationships and causal sequences for any inferred phylogeny.

### Other improvements for phylogenetic analysis using ML

*Fine-grained parallelization for ML analysis*. Many users use MEGA for ML calculations of branch lengths, evolutionary parameters, ancestral states, and divergence times for a given phylogeny. These calculations can be time-consuming for larger datasets, so MEGA12 now implements fine-grained parallelization to speed up the estimation of likelihood values that are calculated independently for different sites at a given node in the phylogeny for a given set of branch lengths and substitution pattern parameter values. Our tests showed a sub-linear reduction in computational time needed, achieving slightly less than 50% efficiency using four threads compared to a single thread. A larger number of threads could offer slightly higher efficiency depending on the number of sequences, variations, and other data attributes. These sub-linear efficiency trends are explained by the fact that substantial overhead is involved in distributing calculations to different threads. In addition, more than half of the nodes in a phylogeny are terminal nodes at which only a few site configurations exist (four or twenty, except for ambiguous states for DNA or protein sequences, respectively), which are not amenable to significant savings.

#### Generating initial trees for heuristic searches for ML phylogenies

Options for automatically generating the initial tree for ML tree searching have been modified in *MEGA12*. When the default option is used, *MEGA12* first generates two initial candidate trees: a neighbor-joining (NJ) tree and a maximum parsimony (MP) tree. The NJ tree is based on evolutionary distances computed using a one-parameter substitution model for nucleotides and amino acid MSAs. To find the MP tree candidate, *MEGA12* conducts ten heuristic searches, each starting with a randomly generated tree subjected to SPR branch swapping; see (Nei and Kumar 2000) for a description. The one with the minimum tree length is chosen among the ten MP trees. Subsequently, the log-likelihood is computed for this MP tree and the NJ tree using the one-parameter substitution model. The tree with superior log likelihood is selected as the initial tree for branch swapping to find the ML tree.

#### Elimination of computational bottlenecks

Testing and benchmarking ML calculations using increasingly larger data sets revealed bottlenecks in the code that were not apparent when using small datasets. For instance, the initialization step to generate a map of identical site patterns was previously done in a way that was too slow for big datasets. *MEGA12* makes this step orders of magnitude faster using a fast hash table. We also identified many instances of redundant initializations (e.g., site configuration maps) and calculations, which have been refactored to speed up calculations.

#### EP calculation updates

The Evolutionary Probabilities (EP) analysis was introduced in *MEGA11* (Tamura et al. 2021) for estimating Bayesian neutral probabilities of observing alternative alleles in a species contingent on the given species phylogeny and the MSA (Liu et al. 2016). The EP analysis in *MEGA* has been updated so that user-provided times, specified as branch lengths in a Newick tree, can be used instead of times computed using RelTime (Tamura et al. 2012). Users can also select the focal sequence via the *Analysis Preferences* dialog box, which was previously restricted to the first sequence in the MSA. The results displayed for the EP calculation have been updated, and the evolutionary timespan of the base (Kumar, Sanderford, et al. 2012) and focal sequence bases for each site are included in the output CSV file.

### Improvements in the Graphical User Interface

The GUI has been updated extensively with many usability improvements and modifications to keep pace with computer hardware, accessories, and operating system changes.

#### Advancement of Tree Explorer (TE)

TE has been enhanced by adding a quick-access panel on the side toolbar to provide easy access to customization options previously accessible only through the menus (see **Fig. 3b**). Searching tip names has been improved to facilitate visualization and navigation through multiple matches. Users can now easily edit the names and fonts of the tip names in the phylogenetic tree, which can now be displayed with equalized branch lengths in TE or with tip names aligned vertically. Labels for internal nodes and group names can now be edited directly in TE by right-clicking a given node. Clones of the *Tree Explorer* and current results can now be generated, giving users snapshot copies of the current display as formatting and other edits are made to one of the copies. Finally, display settings between trees across tabs in TE have been synchronized to align tree displays visually.

#### Advancement of the Tree Topology Editor

MEGA offers functionality for manual drawing and editing a phylogeny, which can help update an existing tree by adding taxa and rearranging them through drag-and-drop operations. The *Tree Topology Editor* in MEGA 12 features several quality-of-life enhancements for manual editing of phylogenies. Users can now assign branch lengths and node heights, and they can see branch lengths and double-click to edit them on the spot, which would come in handy when Newick trees need to have branch lengths or divergence times for display or further calculations, such as EP analysis. By default, the displayed tree now automatically resizes with the window. Moving branches via drag-drop now provides visual feedback to the user. The taxon name editing text box was updated to make the behavior consistent with similar GUI elements in different operating systems.

#### Data Explorer Updates

Responsiveness of scrolling with large data sets has been improved for the *Sequence Alignment Editor* (SAE), *Sequence Data Explorer* (SDE), and *Distance Data Explorer* (DDE). A taxa name search tool and highlighting of all cells corresponding to the current search match have been added for the DDE. In both the SDE and DDE, the sorting of taxa can now be by name or by distance to the first taxon. The number of base differences between the first sequence and all the remaining sequences are used in SDE. When taxa are grouped, individual taxa can be selected/unselected based on many different options: first of each group, by group size, or group inclusion.

#### Dealing with high-resolution monitors

The user experience was severely impacted on computer monitors with ultra-high resolutions when using MEGA11. Standard graphical components (e.g., buttons, icons, and text) are rendered very small on these very high DPI displays. Also, MEGA’s custom visual components, such as the tree display in TE and text grids in SAE, were variously affected by changes in DPI and resolution settings. The problems were more than aesthetic, causing clickable GUI components to be pushed out of view and unusable in some places. Consequently, we needed to redraw hundreds of icons in multiple resolutions and then program MEGA to automatically select the optimal resolution icon images based on the DPI of the monitor. Furthermore, we have updated all the forms and dialog boxes to auto-adjust the size and placement of components based on the monitor resolution.

##### Additional GUI updates

The *MEGA* GUI contains many custom forms to accommodate diverse analyses, results, and data exploration tools. In *MEGA12*, a *Windows* menu has been added to all the data and result explorers, enabling users to navigate to any other currently active windows quickly. We have also made calculation progress reporting more informative, adding analysis details, calculated parameters, and data statistics. The display of some partial results has been programmed when a user issues a command to terminate long-running processes prematurely but desires to see the results obtained thus far, such as the bootstrap analysis. Finally, we have updated the *Caption Expert* system introduced in *MEGA4* (Tamura et al. 2007) to generate natural language descriptions of the models, methods, and parameters used in analyses. All the captions are updated for brevity and clarity. An example caption is shown for a result from *DrPhylo* in **Figure 4d**.

## Conclusions

We have described numerous major upgrades implemented in *MEGA12*, significantly enhancing its computational efficiency and useability. We expect many phylogenetic analyses using ML methods to finish more quickly than previous versions, which is made possible by developing and implementing heuristics that avoid unnecessary computation during the selection of optimal substitution models and bootstrap tests of phylogeny. These heuristics were tested by analyzing many empirical datasets, and the results suggest that their use will generally produce the same result as this without using the heuristics. In the future, we plan to make *MEGA* even more computationally efficient, particularly for analyzing phylogenomic alignments on desktop computers used by many *MEGA* users.

## Acknowledgments

We thank Dr. Alessandra Lamarca for analyzing data to test the initial tree search in the ML phylogeny construction. This work was supported by a research grant from the National Institutes of Health to SK (R35GM139540-04).

## Data and software availability

All sequence alignments and phylogenetic trees used in this article were obtained from published articles and assembled in a Figshare repository: https://figshare.com/s/0413dd262c2ed9df1bd2. *The MEGA12* beta version can be downloaded from https://www.megasoftware.net/beta_download for use on MS Windows. An application for macOS is in the early testing and hardening phase, which we hope to release soon. Linux releases will follow them. At the time of this article’s publication, the final version of the software packages will be available from https://www.megasoftware.net. The source code will be available from https://github.com/KumarMEGALab/MEGA-source-code, which currently contains MEGA11’s source code.

## Materials and Methods

### Data sets analyzed

Simulated datasets were obtained from a previously published research study (Abadi et al. 2019). These nucleotide multiple sequence alignments were generated with varying sequence lengths, number of sequences, base-frequencies, substitution rates, heterogeneity across sites, and proportion of invariant sites. A total of 24 models of substitutions (six base models and their +I, +G, and +I+G combinations; see main text) were used for simulating the data, with 300 datasets generated for each model scenario. From each model category, we randomly selected ten datasets (out of 300), resulting in a total of 240 simulated datasets for model selection analysis. Sequence counts in these datasets ranged from 4 to 289, which were 186 to 18,171 sites long. We performed bootstrapping with the *Adaptive* option and model selection with the *Filtered* option. Then, we estimated their concordance by comparing results from standard bootstrap and model selection with the *Full* option, respectively.

We also analyzed amino acid sequence datasets, generated from a concatenated MSA (*Chloroplast_Martin*.*meg*). The dataset is an example in *MEGA12* (Adachi et al. 2000; Tamura et al. 2021). 45 MSAs were generated using the protein domain boundaries from the concatenated alignment. We performed the model selection analyses for these protein domains and compared the results of the *Full* and *Filtered* options.

The *DrPhylo* analysis was conducted on a clade within a plant phylogeny derived from a maximum likelihood (ML) analysis of an empirical dataset comprising 520 protein-coding genes from 52 plant species (Boachon et al. 2018). This clade was selected because of its reported incongruence with the phylogeny inferred from partitioned data analysis (Shen et al. 2021). The ML tree and multiple sequence alignments (MSAs) for each protein-coding gene were obtained from Boachon et al. (Boachon et al. 2018).

### Options for analyses conducted

We used *MEGA12* for all analyses to directly compare the impact of certain new features while keeping all other aspects the same because many incremental changes and bug fixes have been made over the three years since *MEGA11* was released. For the model selection analysis, we first selected the best-fit substitution models using the *Full* option, which tested all the substitution models. The best-fit models found in these analyses were used as ground truth for model selection results obtained using the *Filtered* option. Default BIC and AICc thresholds of 5 were used in these analyses. For the bootstrap analysis, the *Standard* option in *MEGA12* was used with 500 replicates, and the *Adaptive* option was used with a default SE threshold of 5%.

Model selection and bootstrap analyses were conducted using the command-line version of *MEGA12* (“megacc”) with a single thread for direct comparisons. *DrPhylo* analysis was conducted using *MEGA12*’s GUI. In all these analyses, we used a 64-bit desktop computer with eight logical processors (3.36 GHz) and 64 GB of system memory running the Windows 10 operating system.

## Supplementary Information

**Supplementary Figure S1.**
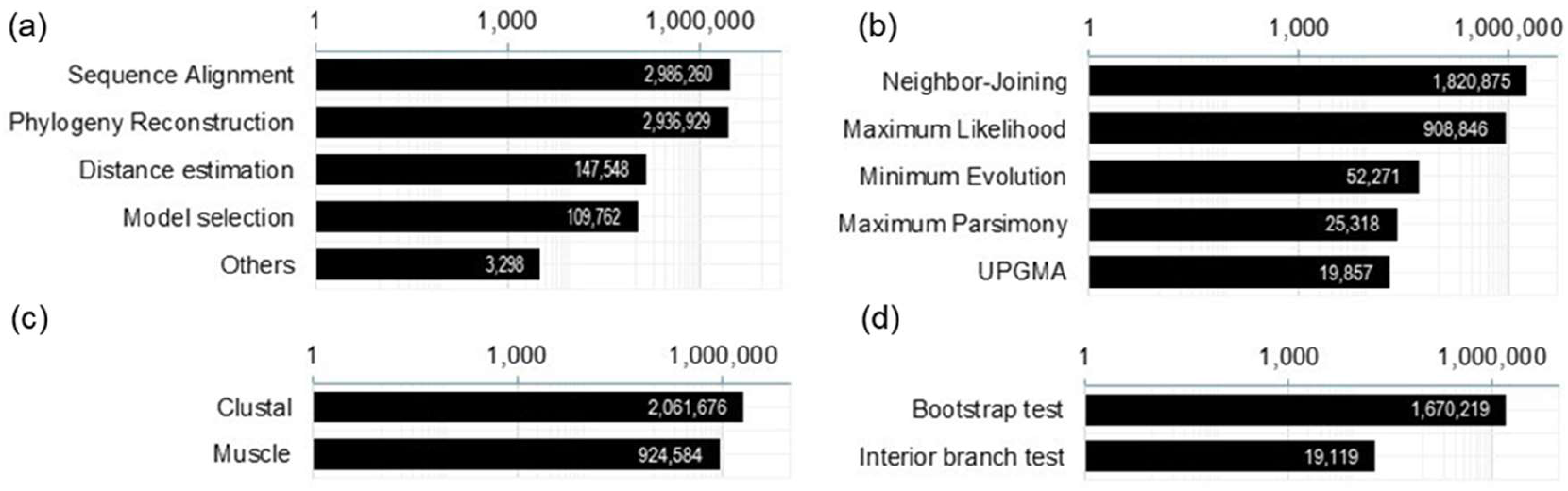
Usage of *MEGA* for various analyses from January 2023 to September 2024. (***a***) Types of analyses. (***b***) Methods of phylogenetic inference. (***c***) Sequence alignment methods. (***d***) Testing of phylogeny. Trends shown are based on data collected using an in-built system to gather anonymous usage data from users who allow this data collection. If a user opts to share their usage data, some versions of *MEGA* save a report of the choices made in the *Analysis Preferences* dialog box. No information about the datasets analyzed is collected, nor is personal or computer information identified. This system is only contained in the GUI version of *MEGA* for the MS *Windows* operating systems, and only a tiny fraction of users permitted data collection. So, these counts are likely to be substantial underestimates of the actual counts of analyses conducted.

**Supplementary Figure S2.**
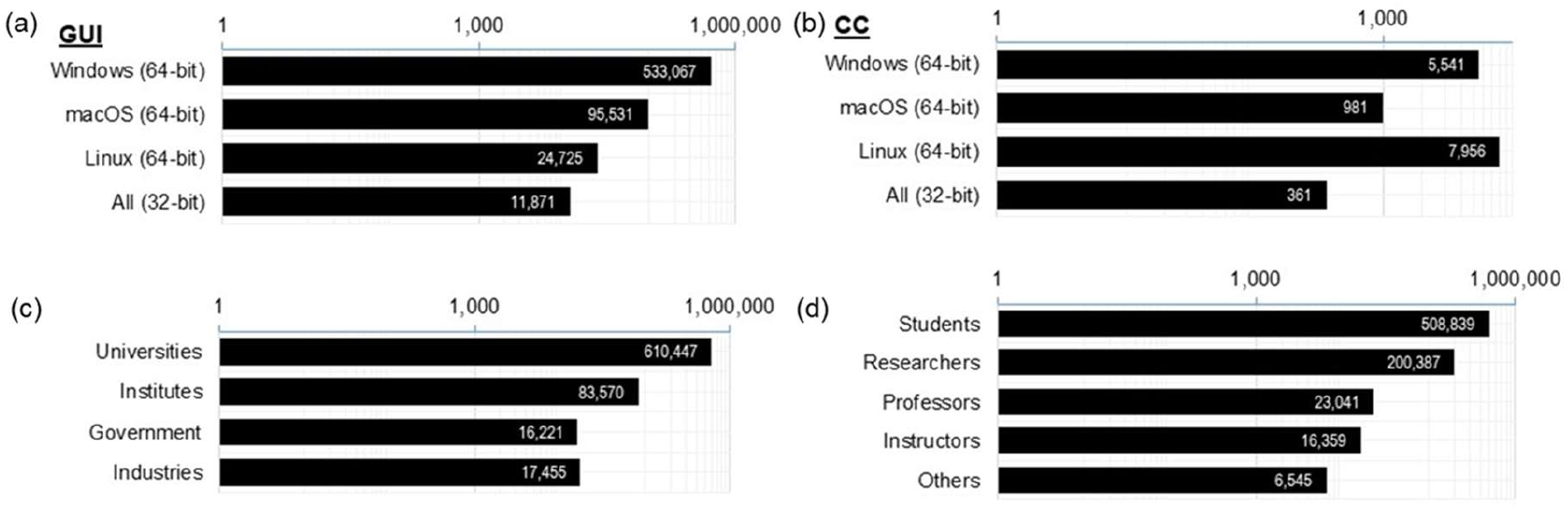
Based on the information received when downloading, downloads, and users of MEGAX and MEGA11 from January 2023 to September 2024. Downloads of (***a***) GUI versions and (***b***) Command-line [CC] versions. (***c***) Types of institutions. (***d***) types of users. Data from Debian, RedHat, and other distributions of Linux are pooled together.

## References

Abadi S, Azouri D, Pupko T, Mayrose I. 2019. Model selection may not be a mandatory step for phylogeny reconstruction. Nat. Commun. 10:1–11.

Adachi J, Waddell PJ, Martin W, Hasegawa M. 2000. Plastid genome phylogeny and a model of amino acid substitution for proteins encoded by chloroplast DNA. J. Mol. Evol. 50:348–358.

Boachon B, Buell CR, Crisovan E, Dudareva N, Garcia N, Godden G, Henry L, Kamileen MO, Kates HR, Kilgore MB, et al. 2018. Phylogenomic Mining of the Mints Reveals Multiple Mechanisms Contributing to the Evolution of Chemical Diversity in Lamiaceae. Mol. Plant 11:1084–1096.

Edgar RC. 2004. MUSCLE: a multiple sequence alignment method with reduced time and space complexity. BMC Bioinformatics 5:113.

Felsenstein J. 1985. Confidence limits on phylogenies: An approach using the bootstrap. Evolution 39:783–791.

Gadagkar SR, Rosenberg MS, Kumar S. 2005. Inferring species phylogenies from multiple genes: concatenated sequence tree versus consensus gene tree. J. Exp. Zool. B Mol. Dev. Evol. 304:64–74.

Gatesy J, Springer MS. 2014. Phylogenetic analysis at deep timescales: unreliable gene trees, bypassed hidden support, and the coalescence/concatalescence conundrum. Mol. Phylogenet. Evol. 80:231–266.

Hedges SB. 1992. The number of replications needed for accurate estimation of the bootstrap P value in phylogenetic studies. Mol. Biol. Evol. 9:366–369.

Kapli P, Yang Z, Telford MJ. 2020. Phylogenetic tree building in the genomic age. Nat. Rev. Genet. 21:428–444.

Kosakovsky Pond SL, Frost SDW. 2005. Not so different after all: a comparison of methods for detecting amino acid sites under selection. Mol. Biol. Evol. 22:1208–1222.

Kumar S. 2022. Embracing Green Computing in Molecular Phylogenetics. Mol. Biol. Evol. 39:msac043.

Kumar S, Filipski AJ, Battistuzzi FU, Kosakovsky Pond SL, Tamura K. 2012. Statistics and truth in phylogenomics. Mol. Biol. Evol. 29:457–472.

Kumar S, Sanderford M, Gray VE, Ye J, Liu L. 2012. Evolutionary diagnosis method for variants in personal exomes. Nat. Methods 9:855–856.

Kumar S, Sharma S. 2021. Evolutionary Sparse Learning for Phylogenomics. Mol. Biol. Evol. 38:4674–4682.

Kumar S, Tamura K, Nei M. 1994. MEGA: Molecular Evolutionary Genetics Analysis software for microcomputers. Comput. Appl. Biosci. 10:189–191.

Kumar S, Tamura K, Nei M. 2004. MEGA3: Integrated software for Molecular Evolutionary Genetics Analysis and sequence alignment. Briefings in bioinformatics 5:150–163.

Liu L, Tamura K, Sanderford MD, Gray VE, Kumar S. 2016. A molecular evolutionary reference for the human variome. Mol. Biol. Evol. 33:245–254.

Nei M, Kumar S. 2000. Molecular evolution and phylogenetics. New York: Oxford University Press

Pattengale ND, Alipour M, Bininda-Emonds ORP, Moret BME, Stamatakis A. 2010. How many bootstrap replicates are necessary? J. Comput. Biol. 17:337–354.

Sanderford M, Sharma S, Tamura K, Stecher G, Liu J, Ji S, Ye J, Kumar S. 2024. MyESL: A software for evolutionary sparse learning in molecular phylogenetics and genomics. Submitted [Internet]. Available from: https://kumarlab.net/downloads/papers/SanderfordKumar2024.pdf

Sharma S, Kumar S. 2021. Fast and accurate bootstrap confidence limits on genome-scale phylogenies using little bootstraps. Nat Comput Sci 1:573–577.

Sharma S, Kumar S. 2024. Discovering fragile clades and causal sequences in phylogenomics by evolutionary sparse learning. Mol. Biol. Evol. 41:msae131.

Shen X-X, Steenwyk JL, Rokas A. 2021. Dissecting Incongruence between Concatenation- and Quartet-Based Approaches in Phylogenomic Data. Syst. Biol. 70:997–1014.

Song S, Liu L, Edwards SV, Wu S. 2012. Resolving conflict in eutherian mammal phylogeny using phylogenomics and the multispecies coalescent model. Proc. Natl. Acad. Sci. U. S. A. 109:14942–14947.

Tamura K, Battistuzzi FU, Billing-Ross P, Murillo O, Filipski A, Kumar S. 2012. Estimating divergence times in large molecular phylogenies. Proc. Natl. Acad. Sci. U. S. A. 109:19333–19338.

Tamura K, Dudley J, Nei M, Kumar S. 2007. MEGA4: Molecular Evolutionary Genetics Analysis (MEGA) software version 4.0. Mol. Biol. Evol. 24:1596–1599.

Tamura K, Peterson D, Peterson N, Stecher G, Nei M, Kumar S. 2011. MEGA5: molecular evolutionary genetics analysis using maximum likelihood, evolutionary distance, and maximum parsimony methods. Mol. Biol. Evol. 28:2731–2739.

Tamura K, Stecher G, Kumar S. 2021. MEGA11: Molecular Evolutionary Genetics Analysis Version 11. Mol. Biol. Evol. 38:3022–3027.

Tamura K, Stecher G, Peterson D, Filipski A, Kumar S. 2013. MEGA6: Molecular Evolutionary Genetics Analysis version 6.0. Mol. Biol. Evol. 30:2725–2729.

Warnow T. 2015. Concatenation Analyses in the Presence of Incomplete Lineage Sorting. PLoS Curr. 7:10.1371/currents.tol.8d41ac0f13d1abedf4c4a59f5d17b1f7.

Williams TA, Cox CJ, Foster PG, Szöllősi GJ, Embley TM. 2020. Phylogenomics provides robust support for a two-domains tree of life. Nat Ecol Evol 4:138–147.

